# Modulation of the sensitivity to ruxolitinib-mediated JAK2 inhibition by mutationally activated SHP2 exhibits cell context dependency in pre-clinical models of myeloproliferative neoplasms

**DOI:** 10.64898/2026.08.19.744423

**Authors:** Tegan M. Rowsell, Garima Pandey, Lucia Mazzacurati, Narmin E. Amin, Gary W. Reuther

## Abstract

Classic Philadelphia chromosome-negative myeloproliferative neoplasms (MPNs) are hematopoietic stem cell cancers that result in aberrant trilineage myeloid cell proliferation, bone marrow fibrosis, and increased risk of acute myeloid leukemia. MPNs are driven by deregulated activity of the JAK2 kinase, induced by mutations in the *JAK2*, *CALR*, and *MPL* genes, but approved JAK2 inhibitors primarily offer palliative effects, not remission. Cell models that demonstrate MPN oncogene driven JAK2 activity requisite for cell proliferation are important research tools for the development of anti-JAK2 and anti-JAK2 signaling therapeutics for MPN. SET2 and UKE1 cells are two such cell lines, as they express JAK2-V617F, one of the major driving mutations of MPN, and require signaling by JAK2 for their growth and viability. These cell lines are AML cell lines that were derived from patients with a previous diagnosis of MPN before they developed AML. Our previous studies demonstrated that the SHP2 phosphatase may be a therapeutic target for MPNs, and here we report our identification and characterization of an activating point mutation of SHP2 (encoded by the *PTPN11* gene), SHP2-F71L, in UKE1 cells. Given SHP2 functions downstream of JAK2 and mediates JAK2 activation of RAS, we set out to determine the effect of mutational activation of SHP2 on the sensitivity of MPN model cells to JAK2 inhibition. We used CRISPR-Cas9 to edit this mutation in UKE1 cells back to wildtype such that these cells only express wildtype SHP2. These cells exhibited enhanced sensitivity to SHP2 inhibition and, notably, enhanced sensitivity to the JAK2 inhibitor ruxolitinib. This altered sensitivity was reverted by exogenous expression of SHP2-F71L but not SHP2-WT, indicating expression of an activated SHP2 may alter sensitivity to JAK2 inhibition in MPN model cells. We further explored this by genetically editing SET2 cells to express SHP2-F71L but observed no change in SHP2 inhibitor or JAK2 inhibitor sensitivity in cells with a SHP2-F71L encoding allele of *PTPN11*. Using the cytokine dependent BaF3 cell line where deregulation of JAK2 signaling by expression of JAK2-V617F induces cytokine independent transformation that remains dependent on this JAK2 signaling, we observed no effect of the expression of an activated SHP2 mutant on the sensitivity of the growth and viability of these cells to ruxolitinib. Recent studies have demonstrated activation of RAS signaling can antagonize JAK2 inhibition in pre-clinical MPN models, and the presence of RAS pathway mutations associates with patients whose disease advances on ruxolitinib therapy. Such mutations include activating mutations in *PTPN11*, as SHP2 is an upstream activator of RAS signaling. Our results suggest that activating *PTPN11* mutations have the potential to desensitize the effects of JAK2 inhibition therapy in patients undergoing therapy and may be dependent on unknown cell and molecular profile contexts.

## Introduction

Myeloproliferative neoplasms (MPNs), including polycythemia vera, essential thrombocythemia, and myelofibrosis, are blood cancers that are driven by aberrant activation of the JAK2 signaling pathway via mutations in three driving oncogenes, *JAK2*, *CALR*, and *MPL*. These cancers are associated with trilineage myeloproliferation, and bone marrow fibrosis in advanced disease.

MPN patients live with significant constitutional symptoms that decrease quality of life and an increased risk of heart attack, stroke, and their disease developing into acute leukemia that is generally recalcitrant to therapy [1–5]. Hematopoietic stem cell transplant offers curative potential for the few who qualify, and clinical studies continue to show promising results for pegylated interferon-alpha for some MPN patients [6–8]. Given the oncogenic driving pathway of these diseases is via aberrantly elevated JAK2 kinase activity, four JAK2 inhibitors are FDA-approved for patients. These drugs decrease symptomology and improve quality of life, but they do not readily, nor substantially, decrease the number of neoplastic cells in patients or induce remission [9–11]. As such, the development of new targeted therapies, including better JAK2 inhibitors (e.g., mutation specific, type II inhibitors), combination therapies with JAK2 inhibitors, or other agents that can target MPN-driving cells continues to be an area of intense research.

Efforts to improve anti-JAK2 or anti-JAK2 signaling will continue to rely on pre-clinical MPN cellular and in vivo therapeutic models. However, while the oncogenic driving etiology of MPNs is well established, there has been an underlying challenge to model MPNs using cell lines. This is based on the fact these cancers, while hematopoietic stem cell disorders like leukemia, are manifest by aberrant production of mature blood cells during the chronic phase of disease in patients, as opposed to hyperproliferation of immature progenitor blast cells as in acute leukemias. Given this, unlike leukemia cell models there are no true MPN cell lines, per se. The MPN cell lines that are used to study anti-JAK2 therapeutics are utilized because they are dependent on the signaling by an MPN driver mutation, most commonly, JAK2-V617F. These human MPN model cell lines include HEL, SET2, and UKE1, each of which is widely used in MPN preclinical studies. These cells contain the activating V617F point mutation of the JAK2 tyrosine kinase, with HEL and UKE1 cells being homozygous for this mutation, and SET2 cells heterozygous, although the ratio of mutant to wildtype JAK2 gene expression in SET2 cells is believed to be about 4 to 1 due to gene amplification [12–14]. The HEL cell line (and its subclone HEL 92.1.7) is a human erythroleukemia cell line originally derived from a patient who had erythroleukemia following Hodgkin’s lymphoma and no reported history of an MPN [13, 15]. SET2 cells were derived from a patient with acute megakaryoblastic leukemia transformation from essential thrombocythemia [16]. UKE1 cells were also derived from a patient with acute leukemia secondary to essential thrombocythemia [17]. While these cells are dependent on JAK2 signaling for growth and viability, as demonstrated by use of small molecule inhibitors, shRNA, and CRISPR essentiality (note: UKE1 cell CRISPR essentiality is not profiled in the public DepMap database, https://depmap.org/portal), they provide cell models for JAK2-V617F signaling to investigate novel anti-JAK2 therapeutics and mechanisms of resistance [12, 13, 18, 19]. Given these cells are in fact AML cell lines, SET2 and UKE1 cells are also utilized as models for JAK2-V617F-positive post-MPN AML [20, 21].

In addition, the murine cytokine dependent BaF3 cell line, which has been used for decades for oncogenic signaling and targeted therapeutic studies wherein exogenous expression of an oncoprotein induces growth factor independent growth, is also used to model JAK2 signaling emanating from the major MPN drivers. This includes cell line models of JAK2-V617F as well as activating mutations of MPL and CALR, all of which are based on growth factor independent growth induced by expression of the MPN driving signaling protein and thus provide isogenic cell models with the difference being the single oncogenic driving signal [22–27].

We recently described the SHP2 tyrosine phosphatase as a therapeutic target in preclinical models of MPN, with SHP2 inhibitors having potential as a single agent as well as in combination with JAK2 inhibition [28]. Our results demonstrated that JAK2 signals through SHP2 to regulate activation of the RAS/ERK pathway, one of the major mediators of JAK2 signaling [28]. During the course of those studies, we identified an activating SHP2 mutation in UKE1 cells. Herein we describe characterization of this mutation and the potential effect mutationally activated SHP2 can have on altering the sensitivity of MPN cell models to JAK2 inhibition.

## Results

In previous studies, we identified the tyrosine phosphatase SHP2 as a therapeutic target for MPN in pre-clinical models, results that have been confirmed by others [28, 29]. Given activating mutants of SHP2 (encoded by the *PTPN11* gene) are known hematopoietic oncoproteins, we sequenced the coding sequences for SHP2 in the UKE1 and SET2 cell lines to confirm the amino acid integrity of the protein expressed as wildtype SHP2 [30]. However, in UKE1 cells we identified a point mutation that changed phenylalanine 71 to leucine (F71L) at both the genomic (Fig.1A, top) and transcript (not shown) levels, with the sequencing indicating this mutation was heterozygous with wildtype. We confirmed this mutation in a second source of UKE1 cells (not shown), suggesting this mutation was not acquired during culture in our laboratory. We then identified the presence of this SHP2-F71L encoding mutation in UKE1 cells in the Harmonizome database [31–33], and since our initial studies, UKE1 cells have been incorporated into the CCLE/DepMap database of the Broad Institute which also lists this F71L mutation of SHP2 in these cells [18, 19]. This mutation is located in the N-SH2 domain of SHP2, and it has been described, like many other missense mutations in the region of amino acids 60- 76 in the SHP2 N-SH2 domain, as an activating mutation due to the destabilization effect such point mutations have on the auto-inhibited confirmation of the enzyme [30, 34, 35]. SHP2 (*PTPN11*) mutations are uncommon in the chronic stage of MPNs, although their rare presence may be associated with a poor response to the JAK2 inhibitor ruxolitinib and their frequency is elevated significantly (∼7-8%) in patients whose chronic MPN disease has transformed to AML [36–38]. Given UKE1 cells were derived from the leukemic transformation of a patient previously diagnosed with essential thrombocythemia, it is most likely this mutation was acquired during this leukemic transformation.

**Figure 1:**
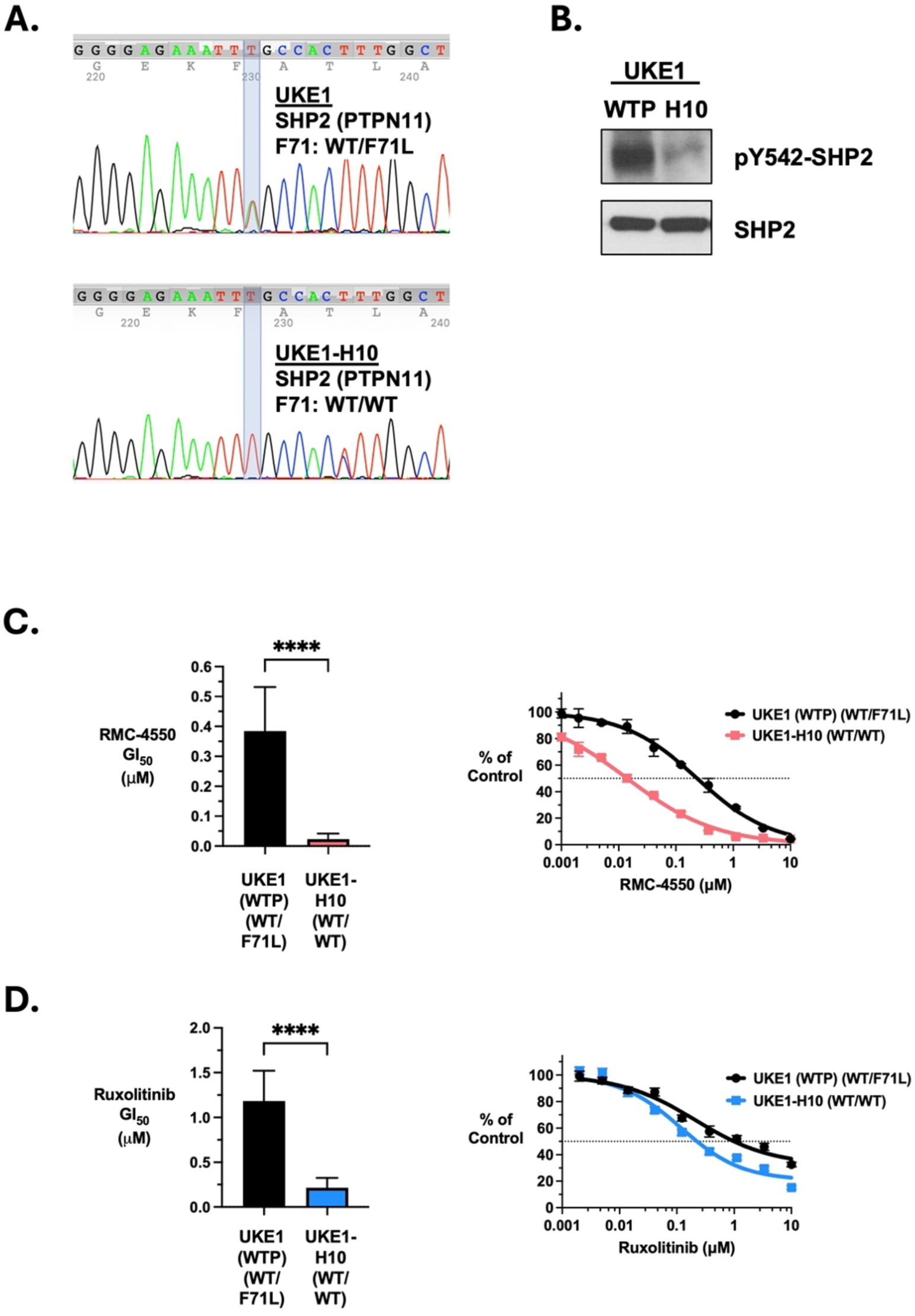
Identification and characterization of a SHP2-F71L-encoding mutation present in the UKE1 MPN model cell line. **A.** Sequencing of cDNA isolated from UKE1 cells indicated the presence of a thymidine to adenosine nucleotide change in the third nucleotide encoding amino acid 71 of SHP2 (not shown). Subsequent PCR and Sanger sequencing analysis of UKE1 cell genomic DNA confirmed this change, which is heterozygous with wildtype (top). Amino acid 71 in wildtype SHP2 is phenylalanine and this base change in one allele of *PTPN11* encodes for leucine, indicating UKE1 cells are heterozygous for *PTPN11* alleles that encode wildtype and SHP2-F71L. This mutation was changed to wildtype (thymidine) by CRISPR-Cas9 and homology-directed repair, generating UKE1-H10 cells, wherein both *PTPN11* alleles encode wildtype SHP2, as shown by PCR and Sanger sequencing of H10 cell genomic DNA (bottom). **B.** Immunoblotting showing pY542-SHP2 and total SHP2 levels in UKE1 (WTP) and H10 cells. WTP cells are UKE1 cells that were put through the CRISPR-Cas9 editing protocol but with Cas9 omitted and thus are unaltered UKE1 cells that express SHP2-F71L from a single *PTPN11* allele. **C.** The GI50 of the SHP2 inhibitor RMC-4550 was determined for UKE1 control and H10 cells. The mean (+/- s.d., n = 8, **** = p value < 0.0001 by t-test) (left) and example concentration-response curves (right) are shown. **D.** The GI50 of the JAK2 inhibitor ruxolitinib was determined for UKE1 control and H10 cells. The mean (+/- s.d., n =10, **** = p value < 0.0001 by t-test) (left) and example concentration-response curves (right) are shown.

SHP2 contributes to activation of RAS signaling by various mechanisms [39–41], and activation of RAS signaling has been shown to antagonize the effects of the JAK2 inhibitor ruxolitinib in preclinical MPN models [42–44]. Given aberrant activity of SHP2 could both enhance RAS activation downstream of deregulated JAK2 signaling in MPN as well as activate RAS independently of JAK2, the presence of the activating mutant SHP2 in UKE1 cells could alter the response of these cells to JAK2 inhibitors, the assessment of which these cells are widely used for. In addition to SHP2 inhibition sensitizing MPN model cells to ruxolitinib, co-targeting of each of MEK and ERK with ruxolitinib in pre-clinical MPN models does as well, suggesting activation of ERK signaling can contribute to JAK2 inhibitor sensitivity [28, 42, 43, 45, 46]. Given this we set out to determine if the F71L activating mutation of SHP2 contributed to the sensitivity of UKE1 cells to JAK2 inhibition. To this end, we utilized CRISPR-Cas9 and homology-directed repair to change the F71L-encoding *PTPN11* mutation back to phenylalanine, the amino acid at this position in the wildtype protein, generating UKE1 cells that were homozygous for wildtype *PTPN11*/SHP2 (Fig. 1A, bottom). These cells, named H10, expressed diminished pY-SHP2 but similar total SHP2 protein compared to control UKE1 cells, in line with expectations of the loss of expression of mutationally activated SHP2 in these cells, and the presence of two wildtype- encoding SHP2 *PTPN11* alleles (Fig. 1B). We and others have reported the sensitivity of pre- clinical MPN models to SHP2 inhibition, including UKE1 cells [28, 29]. Given this, we assessed the relative response of H10 cells compared to UKE1 cells to the SHP2 inhibitor RMC-4550 [47]. H10 cells were remarkably more sensitive to RMC-4550, with an approximate 10-fold shift in the GI50 of this inhibitor compared to control UKE1 cells that went through the genomic editing protocol with transfection of Cas9 only. The GI50 for RMC-4550 was determined to be about 0.35 μM for control cells and 0.02 μM for H10 cells (Fig. 1C). This increased sensitivity is likely due to the fact that activating point mutations of SHP2 that block the protein’s autoinhibitory interactions are less sensitive to allosteric SHP2 inhibitors like RMC-4550 [47, 48]. We then assessed the relative sensitivity of these cells to the JAK2 inhibitor ruxolitinib and determined that H10 cells were more sensitive (about 3 to 4-fold) than control UKE1 cells to this JAK2 inhibitor, suggesting the mutationally active SHP2 present in UKE1 cells may modulate sensitivity to JAK2 inhibition (Fig. 1D).

To further test the potential SHP2-F71L contributes to ruxolitinib sensitivity in UKE1 cells we exogenously expressed SHP2-F71L and SHP2-WT in H10 cells. Exogenous expression of these SHP2 proteins was confirmed by immunoblotting, and this expression was slightly less than endogenous SHP2 protein levels (Fig. 2A). To assess if expression of SHP2-F71L could revert the increased sensitivity of H10 cells to SHP2 inhibition, we determined the GI50 of RMC- 4550 in H10 cells exogenously expressing SHP2 proteins. Cells with exogenous expression of SHP2-F71L were similarly sensitive to SHP2 inhibition by RMC-4550 as parental UKE1 cells, indicating the hypersensitivity of H10 cells to SHP2 inhibition was reverted by the re-expression of SHP2-F71L (Fig. 2B). The high sensitivity of H10 cells to RMC-4550 was not altered by expression of a control vector or expression of a similar level of exogenous SHP2-WT (Fig. 2B). Notably, H10 cells expressing SHP2-F71L exhibited less sensitivity to the JAK2 inhibitor ruxolitinib than H10 cells (Fig. 2C). These cells, in fact, displayed similar sensitivity to ruxolitinib as parental UKE1 cells which express endogenous SHP2-F71L, while H10 cells expressing a control vector or SHP2-WT each displayed similar ruxolitinib GI50s as unmodified H10 cells, indicating the activating SHP2 mutation was responsible for diminishing sensitivity to the JAK2 inhibitor (Fig. 2C).

**Figure 2:**
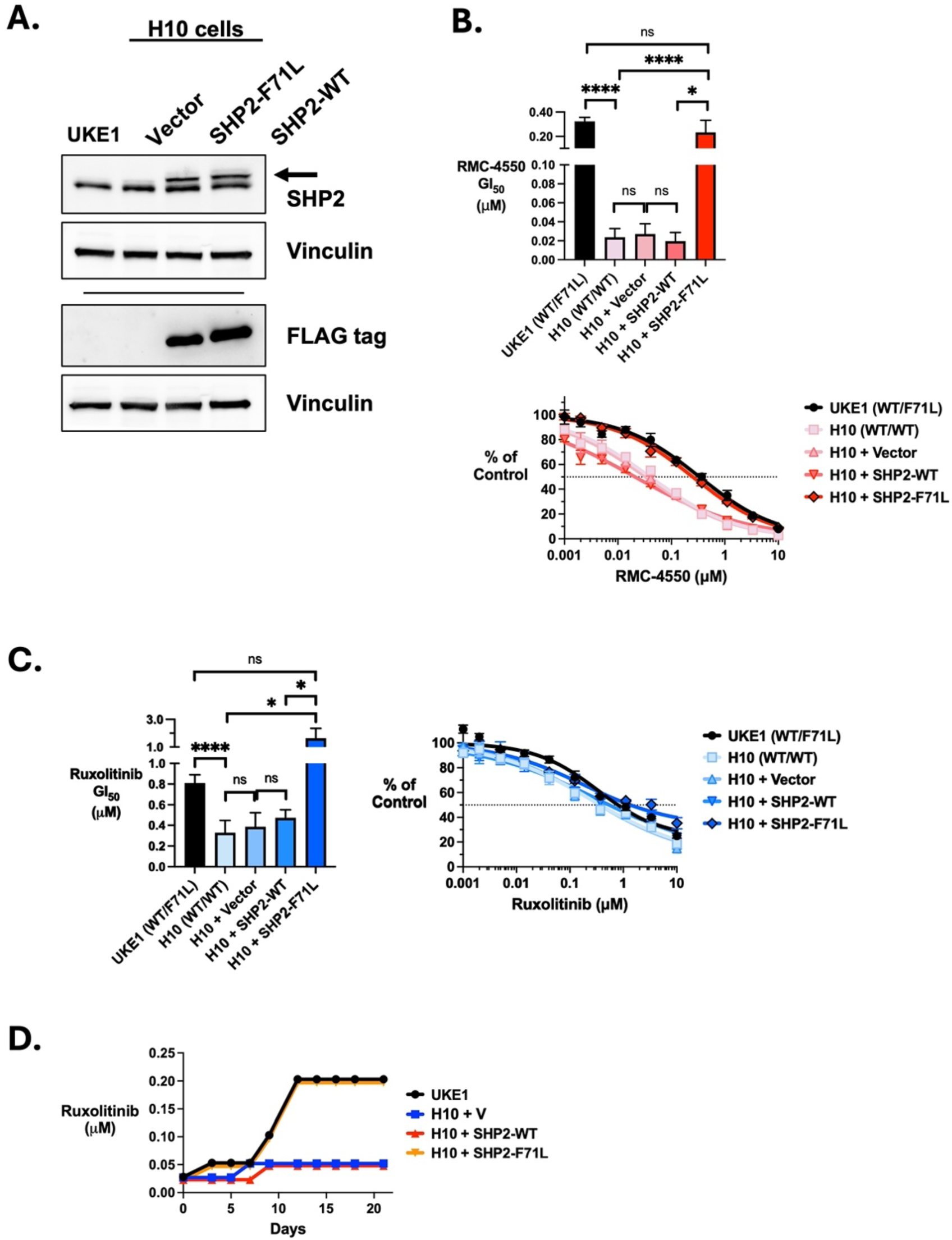
Expression of SHP2-F71L modulates sensitivity of UKE1 cells to both SHP2 and JAK2 inhibition. **A.** UKE1-H10 cells were generated to stably express a control vector, SHP2- F71L-FLAG, or SHP2-WT-FLAG. Immunoblots indicate expression of exogenous SHP2 (arrow) did not exceed endogenous SHP2 levels, as the FLAG tag alters protein mobility to separate it from endogenous SHP2. **B.** The GI50 of the SHP2 inhibitor RMC-4550 was determined for UKE1 and H10 cells, as well as H10 cells expressing a control vector, SHP2-WT, and SHP2- F71L. The mean (+/- s.d., n = at least 3, p values are indicated by **** (p < 0.0001) and * (p < 0.05) by t-test between indicated samples, n.s. = no significant difference) (top) and example concentration-response curves (bottom) are shown. **C.** The GI50 of the JAK2 inhibitor ruxolitinib was determined for UKE1 and H10 cells, as well as H10 cells expressing a control vector, SHP2-WT, and SHP2-F71L. The mean (+/- s.d., n = at least 3, p values are indicated by **** (p < 0.0001) and * (p < 0.05) by t-test between indicated data, n.s. = no significant difference) (left) and example concentration-response curves (right) are shown. **D.** UKE1 and H10 cells expressing a control vector, SHP2-WT, and SHP2-F71L were incubated with DMSO only as well as 25 nM ruxolitinib and total viable cells were determined every 2-3 days. If viable cell numbers of treated cells were 90% of the respective DMSO control treatment, cells were passed back to the initial starting density and the concentration of ruxolitinib was doubled, otherwise the concentration of ruxolitinib was kept the same. Both DMSO and ruxolitinib treated cells were passed back to the original starting cell density at each day cell counting was performed. Shown are the concentrations of ruxolitinib that each cell line was incubated with over time, following this protocol.

While UKE1 cells and H10 cells expressing SHP2-F71L have a higher ruxolitinib GI50 than H10 cells, H10 cells expressing a control vector, or H10 cells expressing SHP2-WT, we wanted to assess if these results from a short term assay translated to an altered ability of these cells to grow in ruxolitinib over time, that is, to develop persistent cell growth in the presence of the JAK2 inhibitor, as we have previously described [49]. After plating cells at a concentration of ruxolitinib that exhibits ∼10% growth inhibition of UKE1 cells (i.e., 0.025 μM, the GI10 at 72 hours), cells were counted every 2-3 days, and their growth was compared to the same cells cultured in DMSO. Cells were replated back to the starting cell density and kept at the same ruxolitinib concentration, unless treated cells grew at a rate of 90% or more of those treated with DMSO, in which case the concentration of ruxolitinib was doubled upon replating. Using this experimental protocol, we determined that UKE1 and H10 cells exogenously expressing SHP2- F71L each adapted to growing in ruxolitinib at the same rate, which was faster than the rate at which H10 cells exogenously expressing SHP2-WT or the control vector adapted (Fig. 2D).

To further investigate if SHP2-F71L expression was sufficient to affect ruxolitinib sensitivity we utilized CRISPR-Cas9 and homology-directed repair to generate SET2 cells to express SHP2- F71L (Fig. 3). Like UKE1 cells, SET2 cells are derived from a post-MPN AML patient and these cells also express JAK2-V617F and their growth and survival are highly sensitive to JAK2 inhibition. While SET2 cells have two *PTPN11* alleles that encode wildtype SHP2 (Fig. 3A, top), the alleles of *PTPN11* in SET2 edited D16 (Fig. 3A, bottom) and E7 (not shown) cells are heterozygous for genes encoding wildtype SHP2 and SHP2-F71L. To assess the relative sensitivity of SET2 cells and D16/E7 cells to JAK2 inhibition we treated each cell line with ruxolitinib. These edited cells that endogenously and heterozygously express SHP2-F71L did not exhibit diminished sensitivity to ruxolitinib (Fig. 3B). Treatment of these cells with the SHP2 inhibitor RMC-4550 did not reveal a clear difference in sensitivity of these cells to SHP2 inhibition, unlike our observations in UKE1 cells. UKE1 and SET2 cells are patient-derived AML cell lines with different genetic profiles.

**Figure 3:**
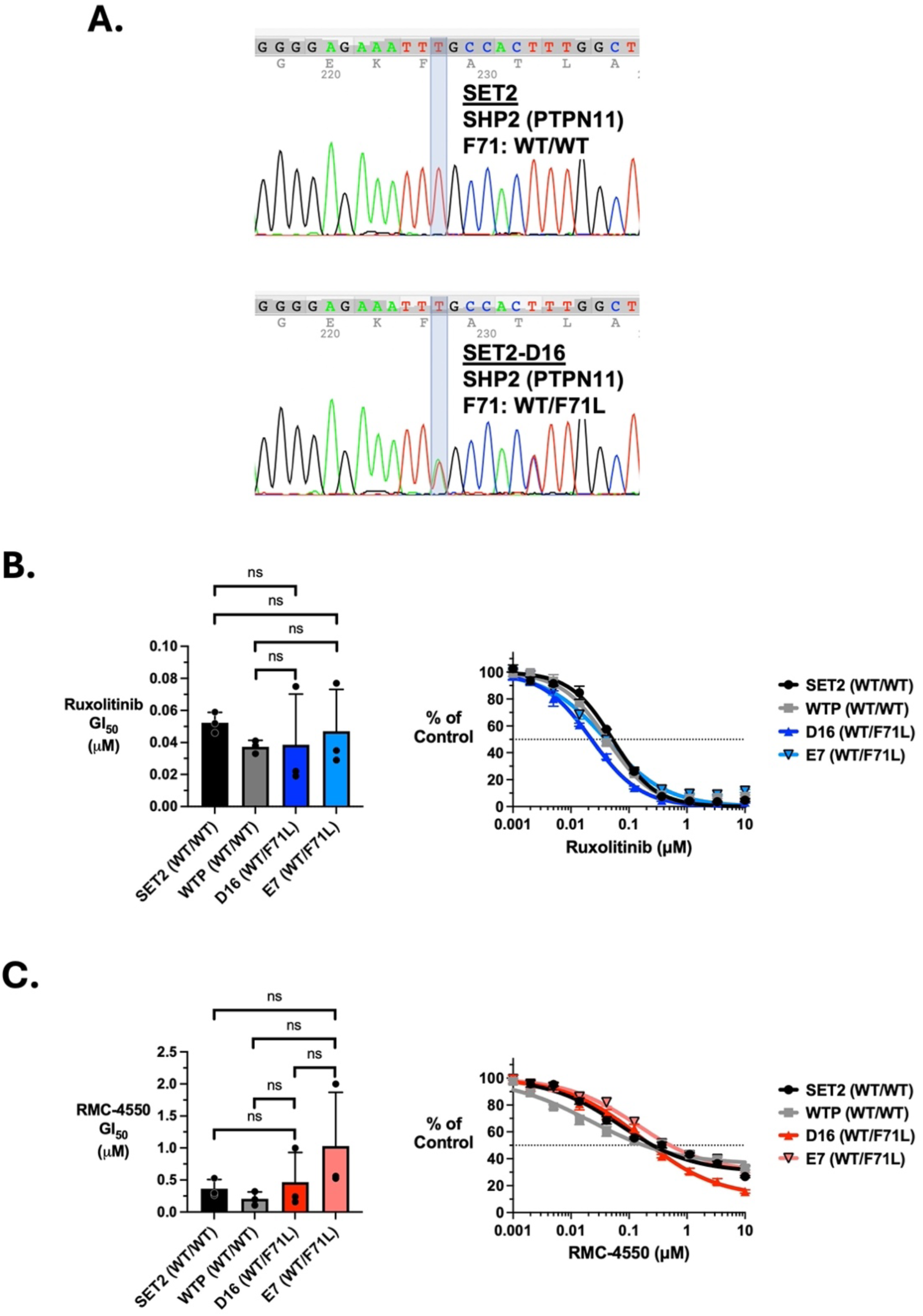
Introduction of a *PTPN11* point mutation that encodes for SHP2-F71L does not alter sensitivity of SET2 cells to SHP2 or JAK2 inhibition. A. The region of *PTPN11* encoding SHP2-F71 was amplified by PCR and submitted for Sanger sequencing, confirming both alleles of *PTPN11* encode for wildtype phenylalanine at amino acid 71 of SHP2 (top). SET2-D16 cells (and E7 cells, not shown) were generated by CRISPR-Cas9 and homology-directed repair to convert the third nucleotide of codon 71 of *PTPN11*/SHP2 from thymidine to adenosine. Confirmation of this mutation was determined by PCR and Sanger sequencing, demonstrating one allele of *PTPN11* in these SET2 cells now encodes for SHP2-F71L (bottom), as in UKE1 cells. **B.** The GI50 of the JAK2 inhibitor ruxolitinib was determined for SET2, control WTP, D16, and E7 cells. The mean (+/- s.d., n = 3 experiments, n.s. = no significant difference by t-test between indicated data) (left) and example concentration-response curves (right) are shown. **C.** The GI50 of the SHP2 inhibitor RMC-4550 was determined for SET2, control WTP, D16, and E7 cells. The mean (+/- s.d., n = 3 experiments, n.s. = no significant difference by t-test between indicated data) (left) and example concentration-response curves (right) are shown.

To expand our studies to a more controlled isogenic cell system that is driven by expression of JAK2-V617F, we utilized the mouse cytokine dependent BaF3 cell line wherein expression of JAK2-V617F along with EpoR induces cytokine independent proliferation [23, 24, 50]. These cells require activated JAK2 signaling induced by JAK2-V617F for maintained proliferation and survival and are widely used in pre-clinical MPN studies to assess anti-JAK2 therapeutics. We expressed wildtype SHP2 or an activated SHP2-E76K mutant, to expand our study beyond SHP2-F71L and assess a well characterized SHP2 mutant, in cytokine independent BaF3-EpoR-JAK2-V617F cells (Fig. 4A) and observed no effect on ruxolitinib sensitivity of this cell model of oncogenic MPN signaling by expression of a mutationally activated SHP2 (Fig. 4B).

**Figure 4:**
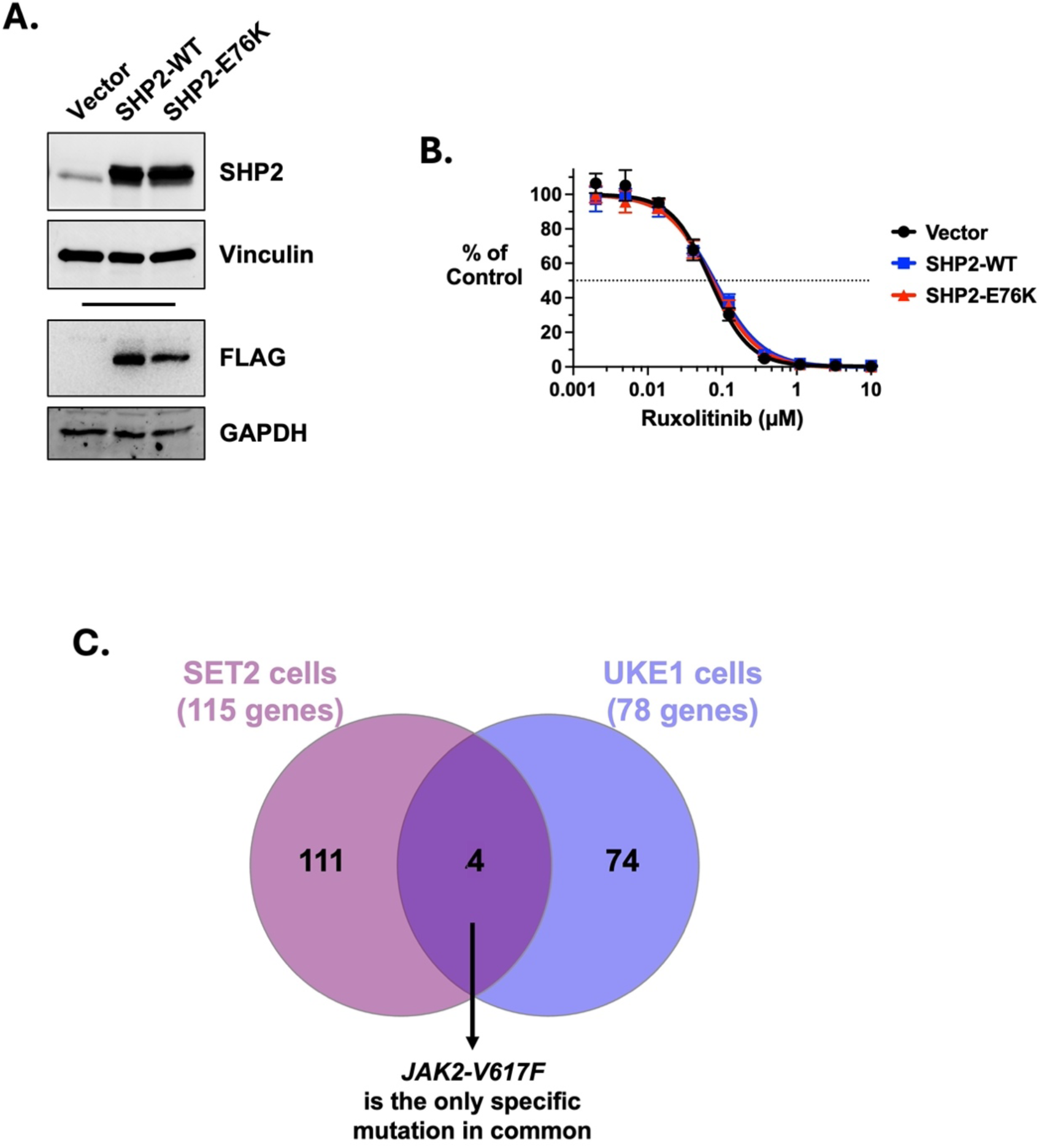
Activated SHP2 expression does not alter ruxolitinib sensitivity to JAK2-V617F- mediated cytokine independent growth of BaF3 cells. **A**. Cytokine independent BaF3-EpoR- JAK2-V617F cells stably expressing a control vector, SHP2-WT-FLAG, and SHP2-E76K-FLAG were generated. Immunoblot analysis showing expression of SHP2-WT and SHP2-E76K in these cells is shown. **B.** The GI50 of ruxolitinib against these cells was determined, with a representative experiment shown. **SET2 and UKE1 cells have dissimilar mutational profiles. C.** Comparison of the gene mutational profiles of SET2 and UKE1 cells as described in the text are shown, with only one specific mutation, that encoding JAK2-V617F, being identical between these two MPN signaling model cell lines (numbers indicate numbers of genes mutated based on the DepMap database) [18, 19].

## Discussion

The inability of JAK2 inhibition to readily induce remission in patients is a clinical bottleneck of current driver targeted therapeutic approaches for MPN patients. Drug resistance-inducing mutations in JAK2 are not evident in patients who lose responsiveness to JAK2 inhibition, and thus are not responsible for the loss of efficacy of JAK2 inhibitors in patients [51]. The persistent survival of disease-driving cells during JAK2 inhibitor therapy has been investigated in pre- clinical MPN models where both JAK2-dependent and independent mechanisms of reactivation of JAK2 signaling pathways have been described. JAK2 inhibitor persistence was first described as being due to inhibitor induced heterodimerization between JAK2 and JAK1 or Tyk2, where this heterodimerization led to reactivation of JAK2 signaling [52]. How this mechanism works remains unclear, especially given that ruxolitinib is a potent JAK1 inhibitor as well. In addition, JAK2 independent pathways that induce activation of ERK signaling have been described using MPN mouse models, where JAK2 inhibitor therapy induces growth factor receptor tyrosine kinase signaling, leading to activation of ERK in a JAK2 independent manner [43]. Thus, this may induce cell survival signals, via activation of receptor tyrosine kinase signaling including RAS/ERK, that overcome the effect of JAK2 inhibition. It has also been shown that expression of activated RAS and MEK proteins antagonize the effects of JAK2 inhibition in JAK2-V617F- driven MPN model cells [44]. Some of these and other reports demonstrate that inhibition of MEK and ERK with direct kinase inhibitors can improve therapeutic responses in pre-clinical models [43, 45, 46]. Likewise, our previous studies demonstrated that the SHP2 phosphatase regulates activation of RAS activation, and subsequently ERK signaling, in MPN models and may be a target to improve JAK2 inhibitor-based therapies [28]. The ERK regulator DUSP6 and downstream ERK target RSK1 may also contribute to JAK2 inhibitor resistance, further highlighting the importance of the RAS/ERK pathway in antagonizing effects of JAK2 inhibition [53]. Sequencing efforts of cells from myelofibrosis patients on ruxolitinib have determined that RAS pathway mutations become associated with ruxolitinib therapy failure and advancing disease, further suggesting this pathway may contribute to the dynamics of JAK2 inhibitor responses in patients [54–60]. Interestingly, a recent study demonstrated JAK2 inhibition can actually enhance oncogenic RAS signaling in myelofibrosis, making the dynamics between JAK2 and RAS signaling more complex yet important to understand [56].

We identified a known activating mutation in SHP2, SHP2-F71L, in the UKE1 cell line that is widely used as a cell model for MPN signaling, specifically JAK2-V617F signaling, and MPN therapeutic studies. UKE1 cells were derived from a patient previously diagnosed with essential thrombocythemia whose MPN had transformed to AML [17]. While mutations in the gene that encodes SHP2, *PTPN11*, are rarely present in MPN patients, the frequency of activating mutations in *PTPN11* is elevated in post-MPN AML (∼7-8%) [36–38]. Thus, this SHP2 mutation in UKE1 cells is likely to have been associated with the patient’s MPN transforming to AML. This mutation is identified in UKE1 cells in two public databases, including DepMap/CCLE, which recently incorporated UKE1 cells [18, 19]. In addition to AML (∼5%), including therapy-related AML (∼15%), activating mutations in *PTPN11*/SHP2 are frequently found in juvenile myelomonocytic leukemia (∼35%) [61, 62].

Our use of genetic editing to generate UKE1 cells with *PTPN11* alleles that both encode for wildtype SHP2 suggested that the expression of SHP2-F71L in UKE1 cells significantly affected the response of these cells to SHP2 inhibition by RMC-4550, which agrees with the fact that activating SHP2 mutations are less sensitive to allosteric SHP2 inhibitors [47, 48]. Reverting the activating SHP2-encoding gene to wildtype increased the sensitivity of the growth of the cells to ruxolitinib suggesting expression of SHP2-F71L may be able to regulate cellular responses to JAK2 inhibition. Importantly, the effects of this genetic editing, the increased sensitivity to SHP2 inhibition and the increased sensitivity to ruxolitinib were reversed by exogenous expression of SHP2-F71L, but not SHP2-WT, indicating mutationally activated SHP2 can modulate JAK2 inhibitor sensitivity.

We used a similar genetic editing approach to generate MPN model SET2 cells to contain one allele of *PTPN11* that encodes SHP2-F71L and determined that this did not alter the sensitivity of these cells to either SHP2 inhibition by RMC-4550 or to JAK2 inhibition by ruxolitinib. While UKE1 and SET2 cells are patient-derived AML cell lines that both express the JAK2-V617F mutation and whose growth and viability are sensitive to JAK2 inhibitors, these cell models of course differ at their genetic level. For example, while SET2 cells are heterozygous for JAK2- V617F and homozygous for wildtype SHP2, UKE1 cells are homozygous for JAK2-V617F and contain one *PTPN11* allele that encodes for SHP2-F71L. While it isn’t possible to tell if the SHP2-F71L mutation in UKE1 cells was a driver of the AML transformation in the patient these cells were derived from, or if it was eventually selected for by further clonal evolution over time, it is certainly possible that this evolution positively supported signaling (e.g., elevated RAS signaling) by the mutationally activated SHP2. This evolution leading to the isolation of UKE1 cells may have created an intrinsic cellular environment in which JAK2 inhibitor sensitivity is modulated by mutationally active SHP2. This may be unlike in SET2 cells, which of course had its unique evolutionary development that may have led to the sensitivity of these cells to JAK2 inhibition not being readily influenced by an activated SHP2. Utilizing the DepMap database mutation profile (including single nucleotide variants, point mutations, insertions, deletions, and substitutions) of each of these cell lines, which lists mutations in 115 genes in SET2 cells and in 78 genes in UKE1 cells, only four genes are found in common to be mutated in these two cell lines, with only the specific JAK2-V617F-encoding mutation being a common mutation between them (Figure 4C) [18, 19].

The ability of mutationally active SHP2 to alter sensitivity to ruxolitinib may differ by specific SHP2 activating mutations and cell context dependencies that define differential strength of SHP2-mediated RAS signaling. Mutational activation of RAS/ERK signaling driven by mutant RAS may induce stronger deregulated signaling than expression of point-mutationally activated SHP2, which while also leading to RAS activation, leaves this activation in a state that can still be controlled by negative regulatory mechanisms (e.g., the intrinsic GAP activity of RAS, and RAS-GAP protein activity). Given activation of the RAS/ERK pathway is highly regulated by feedback inhibitory signaling loops, it is possible mutationally activated SHP2 may not associate with JAK2 inhibitor sensitivity, even though wildtype SHP2 functions to regulate RAS activation downstream of JAK2.

It is possible post-MPN AML patients with activating *PTPN11*/SHP2 mutations will be less responsive to JAK2 inhibition, in line with multiple lines of evidence indicating activation of the RAS/ERK pathway may contribute to inefficacy of JAK2 inhibitors [28, 42–46, 56]. However, our results suggest this may be dependent on specific cellular and genetic contexts, as expression of an activated SHP2-F71L renders UKE1 cells, but not SET2 cells, less sensitive to JAK2 inhibition. This also renders UKE1 but not SET2 cells less sensitive to SHP2 inhibition suggesting UKE1 cells may be more dependent on, or are able to support the elevated signaling of, activated SHP2. The selection of RAS mutations in patients on ruxolitinib therapy has been described as a paradoxical reciprocal activation, where RAS activation downregulates JAK2 signaling, and vice versa, leaving cells with RAS mutations less reliant on aberrant JAK2 signaling and thus less responsive to JAK2 inhibition [56]. Whether or not the same is true of activating *PTPN11*/SHP2 mutations that constitutively signal toward RAS activation is unknown.

In summary, our results indicate that point mutations in the *PTPN11* gene that result in expression of constitutively active SHP2 protein have the potential to modulate sensitivity to JAK2 inhibition in a cell intrinsic manner. However, this may only be true in certain cellular and genetic contexts. While beyond the scope of our study to assess, extrinsic signals from inflammatory cytokines and the bone marrow microenvironment may be modulated by the presence of mutationally activated SHP2, and together with effects on JAK2 and RAS signaling pathways this may also alter JAK2 inhibitor sensitivity.

## Methods

### Cell culture

UKE1 cells were obtained from Coriell Institute and SET2 cells were obtained from DSMZ (German Collection of Microorganisms and Cell Cultures GmbH). Cell authenticity was confirmed by short tandem repeat profiling. UKE1 cells were cultured in RPMI supplemented with 10% fetal bovine serum (FBS), 5% horse serum, 1 mM hydrocortisone, and penicillin/streptomycin. SET2 cells were cultured in RPMI supplemented with 10% FBS and penicillin/streptomycin. BaF3-EpoR-JAK2-V617F cells were previously described and cultured in RPMI supplemented with 10% FBS and penicillin/streptomycin [63, 64].

### GI50 Concentration Determination

The concentrations of inhibitors that reduce viable cells to 50% of an untreated control (GI50) were determined using Cell-Titer Glo 2.0 (Promega Corporation) following incubation of cells with a range of inhibitor concentrations and analysis with SoftMax Pro 7.1 (Molecular Devices, LLC), and Prism 11 (GraphPad Software, LLC) was used for graphing and statistical analyses, as described previously [28]. GI50 assessment in UKE1 and SET2 cells was after 72 hours of inhibitor incubation, while the GI50 assessment for BaF3-EpoR-JAK2-V617F cells was after 48 hours. The development of ruxolitinib persistent growth was done as described previously [49], and in the legend of Figure 2.

### CRISPR-Cas9 editing

UKE1 cells were edited to generate UKE1 cells that lack the F71L- encoding mutation of SHP2 in *PTPN11*, and SET2 cells were edited to generate SET2 cells that contain an F71L-encoding mutation of SHP2 in *PTPN11* by CRISPR-Cas9 and homology-directed repair (Advanced Cells/Synthego Corporation, now EditCo Bio, Inc.). To revert the SHP2-L71 codon to F71 (wildtype) in the L71-encoding *PTPN11* allele in UKE1 cells, the guide RNA sequence used was AGGGGAGAAA<u>UUA</u>GCCACUU and the donor DNA sequence was ACACTGGTGATTACTATGACCTGTATGGAGGGGAGAAA<u>TTT</u>GCCAC<u>C</u>TTGGCTGAGTTGGT CCAGTATTACATGGAACATC. The SHP2-L71 codon is underlined in the guide RNA sequence, and the SHP2-F71 codon is underlined in the donor DNA sequence. A silent point mutation (underlined) was introduced in the donor sequence to change the PAM site for the guide RNA to reduce cutting post-editing, and this nucleotide change can be seen in the sequencing analysis in Figure 1. To mutate the SHP2-F71 (wildtype) codon to L71 in SET2 cells, the guide RNA sequence used was AGGGGAGAAA<u>UUU</u>GCCACUU and the donor DNA sequence was ACACTGGTGATTACTATGACCTGTATGGAGGGGAGAAA<u>TTA</u>GCCAC<u>C</u>TTGGCTGAGTTGGT CCAGTATTACATGGAACATC. The SHP2-F71 codon is underlined in the guide RNA sequence, and the SHP2-L71 codon is underlined in the donor DNA sequence. A silent point mutation (underlined) was introduced in the donor sequence to change the PAM site for the guide RNA to reduce cutting post-editing, and this nucleotide change can be seen in the sequencing analysis in Figure 3. PCR primers used to amplify the edited region from genomic DNA were: Forward (5’-3’): TGTTGAGTTGGTTGACATGTGG and Reverse (5’-3’): AGCAGCAGACTTTGTGGTCA. PCR products were primed for Sanger Sequencing with the Forward primer (GENEWIZ from Azenta Life Sciences).

### Immunoblotting, antibodies, and inhibitors

Cells were lysed in RIPA buffer containing protease and phosphatase inhibitors (9806, Cell Signaling Technology, Inc.), protein concentrations were determined by BCA assay (Thermo Fisher Scientific, Inc.), and equal lysate protein was analyzed by immunoblotting, as previously described [28]. Antibodies used include those that detect total SHP2 (#7384) and Vinculin (#73614) (Santa Cruz Biotechnology), and pSHP2 (Y542) (#3751), DYKDDDDK Tag (#2368) (to detect FLAG), and GAPDH (#5174) (Cell Signaling Technology). Primary antibodies were detected using secondary antibodies conjugated to horse radish peroxidase for chemiluminescent detection (Thermo Fisher Scientific, Inc.) and StarBright Blue 520 and StarBright Blue 700 (Bio-Rad Laboratories, Inc.) for fluorescent detection, and blots were imaged using a Chemidoc™ MP imaging system (Bio-Rad Laboratories, Inc.). RMC-4550 was obtained from Revolution Medicines, Inc.

### Exogenous SHP2 expression

Wildtype human SHP2 coding sequence was cloned into pHR- SIN-CSGW [65] utilizing In-Fusion® cloning (Takara Bio Inc.) to create pHR-hSHP2-P2A-eGFP which is designed to express human SHP2 and eGFP from a single transcript that also encodes a FLAG tag followed by a P2A self-cleaving peptide both in frame downstream of SHP2 and upstream of eGFP coding sequences. Site-directed mutagenesis was performed using PrimeStar HS (Takara Bio, Inc.) to generate point mutations in the SHP2 coding sequence of this plasmid. DNA sequences were confirmed by Sanger sequencing (GENEWIZ from Azenta Life Sciences) and Whole Plasmid Sequencing (Oxford Nanopore, R10.4.1) (Plasmidsaurus Inc.). Lentivirus was generated by co-transfection of these plasmids with psPAX2, which was a gift from Didier Trono (Addgene plasmid # 12260; http://n2t.net/addgene:12260; RRID:Addgene_12260) and a plasmid that encodes for the VSV-G envelope protein into 293T cells using TransIT-VirusGEN® Transfection Reagent (Mirus Bio) following the manufacturer’s protocol for the transfection reagent. Two days after transfection the medium from transfected cells was collected, filtered, mixed with equal volume of target cell growth medium containing polybrene and used to infect target cells by spinfection at 1,800xg for 90 minutes. Cells were resuspended in fresh growth medium and expanded, resulting in cell populations that were approximately 10% to 30% positive for GFP (assessed using a BD FACSCanto™ 10 Color, BD Biosciences), which were sorted for GFP using a MACSQuant® Tyto® Cell Sorter (Miltenyi Biotec).

## Acknowledgments

This work was supported by the National Institutes of Health grant R01HL151579 from the National, Heart, Lung, and Blood Institute. This work has been supported in part by the Flow Cytometry Core and the Analytic Microscopy Core at the H. Lee Moffitt Cancer Center & Research Institute, a comprehensive cancer center designated by the National Cancer Institute and funded in part by Moffitt’s Cancer Center Support Grant (P30-CA076292).

## References

1. Constantinescu SN, Vainchenker W, Levy G, Papadopoulos N. Functional Consequences of Mutations in Myeloproliferative Neoplasms. Hemasphere. 2021;5:e578.

2. Levine RL, Pardanani A, Tefferi A, Gilliland DG. Role of JAK2 in the pathogenesis and therapy of myeloproliferative disorders. Nat Rev Cancer. 2007;7:673–83.

3. Spivak JL. Myeloproliferative Neoplasms. N Engl J Med. 2017;376:2168–81.

4. Tefferi A, Pardanani A. Myeloproliferative Neoplasms: A Contemporary Review. JAMA Oncol. 2015;1:97–105.

5. Vainchenker W, Kralovics R. Genetic basis and molecular pathophysiology of classical myeloproliferative neoplasms. Blood. 2017;129:667–79.

6. Bose P, Xiao Z, Hasselbalch HC, Prchal JT, Duan M, Yacoub A, et al. Highlights from MPN Asia 2025: Advances in Molecular Pathogenesis and Therapeutic Strategies in Myeloproliferative Neoplasms. Curr Hematol Malig Rep. 2025;20:9.

7. Metzger M, Mascarenhas J. Interferon alpha in myeloproliferative neoplasms: evidence and practical considerations for clinical care. Leuk Lymphoma. 2026;67:489–511.

8. Silver RT, Hasselbalch HC. A paradigm shift in the treatment of patients with polycythemia vera. The initial early use of recombinant interferon-alpha. Leukemia. 2026;40:1122–36.

9. Constantinescu SN, Vainchenker W, Pecquet C. Next-generation JAK inhibitors in the treatment of myeloproliferative neoplasms. Blood. 2026;147:1255–66.

10. How J, Garcia JS, Mullally A. Biology and therapeutic targeting of molecular mechanisms in MPNs. Blood. 2023;141:1922–33.

11. Loscocco GG, Guglielmelli P. Targeted Therapies in Myelofibrosis: Present Landscape, Ongoing Studies, and Future Perspectives. Am J Hematol. 2025;100 Suppl 4:30–50.

12. Jedidi A, Marty C, Oligo C, Jeanson-Leh L, Ribeil JA, Casadevall N, et al. Selective reduction of JAK2V617F-dependent cell growth by siRNA/shRNA and its reversal by cytokines. Blood. 2009;114:1842–51.

13. Quentmeier H, MacLeod RA, Zaborski M, Drexler HG. JAK2 V617F tyrosine kinase mutation in cell lines derived from myeloproliferative disorders. Leukemia. 2006;20:471– 6.

14. Jones AV, Bunyan DJ, Cross NC. No evidence for amplification of V617F JAK2 in myeloproliferative disorders. Leukemia. 2007;21:2561–3.

15. Martin P, Papayannopoulou T. HEL cells: a new human erythroleukemia cell line with spontaneous and induced globin expression. Science. 1982;216:1233–5.

16. Uozumi K, Otsuka M, Ohno N, Moriyama T, Suzuki S, Shimotakahara S, et al. Establishment and characterization of a new human megakaryoblastic cell line (SET-2) that spontaneously matures to megakaryocytes and produces platelet-like particles. Leukemia. 2000;14:142–52.

17. Fiedler W, Henke RP, Ergun S, Schumacher U, Gehling UM, Vohwinkel G, et al. Derivation of a new hematopoietic cell line with endothelial features from a patient with transformed myeloproliferative syndrome: a case report. Cancer. 2000;88:344–51.

18. Arafeh R, Shibue T, Dempster JM, Hahn WC, Vazquez F. The present and future of the Cancer Dependency Map. Nat Rev Cancer. 2025;25:59–73.

19. DepMap BDPQDdo. DepMap, Broad. DepMap Public 26Q1. Dataset. depmap.org. 2026.

20. Fiskus W, Mill CP, Bose P, Masarova L, Pemmaraju N, Dunbar A, et al. Preclinical efficacy of CDK7 inhibitor-based combinations against myeloproliferative neoplasms transformed to AML. Blood. 2025;145:612–24.

21. Wang Z, Skwarska A, Poigaialwar G, Chaudhry S, Rodriguez-Meira A, Sui P, et al. Efficacy of a novel BCL-xL degrader, DT2216, in preclinical models of JAK2-mutated post-MPN AML. Blood. 2025;146:341–55.

22. Daley GQ, Baltimore D. Transformation of an interleukin 3-dependent hematopoietic cell line by the chronic myelogenous leukemia-specific P210bcr/abl protein. Proc Natl Acad Sci U S A. 1988;85:9312–6.

23. James C, Ugo V, Le Couedic JP, Staerk J, Delhommeau F, Lacout C, et al. A unique clonal JAK2 mutation leading to constitutive signalling causes polycythaemia vera. Nature. 2005;434:1144–8.

24. Levine RL, Wadleigh M, Cools J, Ebert BL, Wernig G, Huntly BJ, et al. Activating mutation in the tyrosine kinase JAK2 in polycythemia vera, essential thrombocythemia, and myeloid metaplasia with myelofibrosis. Cancer Cell. 2005;7:387–97.

25. Pikman Y, Lee BH, Mercher T, McDowell E, Ebert BL, Gozo M, et al. MPLW515L is a novel somatic activating mutation in myelofibrosis with myeloid metaplasia. PLoS Med. 2006;3:e270.

26. Klampfl T, Gisslinger H, Harutyunyan AS, Nivarthi H, Rumi E, Milosevic JD, et al. Somatic mutations of calreticulin in myeloproliferative neoplasms. N Engl J Med. 2013;369:2379–90.

27. Nangalia J, Massie CE, Baxter EJ, Nice FL, Gundem G, Wedge DC, et al. Somatic CALR mutations in myeloproliferative neoplasms with nonmutated JAK2. N Engl J Med. 2013;369:2391–405.

28. Pandey G, Mazzacurati L, Rowsell TM, Horvat NP, Amin NE, Zhang G, et al. SHP2 inhibition displays efficacy as a monotherapy and in combination with JAK2 inhibition in preclinical models of myeloproliferative neoplasms. Am J Hematol. 2024;99:1040–55.

29. Mohi G, Yang Y, Sayem MA, Le BT, Ather F, Dutta A, et al. Genetic Deletion or Pharmacologic Inhibition of PTPN11 Impedes the Development and Progression of Myeloproliferative Neoplasms Induced By JAK2V617F and MPLW515L Mutants. Blood. 2023;142:739–39.

30. Chan G, Kalaitzidis D, Neel BG. The tyrosine phosphatase Shp2 (PTPN11) in cancer. Cancer Metastasis Rev. 2008;27:179–92.

31. Rouillard AD, Gundersen GW, Fernandez NF, Wang Z, Monteiro CD, McDermott MG, et al. The harmonizome: a collection of processed datasets gathered to serve and mine knowledge about genes and proteins. Database (Oxford). 2016;2016.

32. Diamant I, Clarke DJB, Evangelista JE, Lingam N, Ma’ayan A. Harmonizome 3.0: integrated knowledge about genes and proteins from diverse multi-omics resources. Nucleic Acids Res. 2025;53:D1016–D28.

33. Klijn C, Durinck S, Stawiski EW, Haverty PM, Jiang Z, Liu H, et al. A comprehensive transcriptional portrait of human cancer cell lines. Nat Biotechnol. 2015;33:306–12.

34. Bentires-Alj M, Paez JG, David FS, Keilhack H, Halmos B, Naoki K, et al. Activating mutations of the noonan syndrome-associated SHP2/PTPN11 gene in human solid tumors and adult acute myelogenous leukemia. Cancer Res. 2004;64:8816–20.

35. Tartaglia M, Martinelli S, Stella L, Bocchinfuso G, Flex E, Cordeddu V, et al. Diversity and functional consequences of germline and somatic PTPN11 mutations in human disease. Am J Hum Genet. 2006;78:279–90.

36. Lasho TL, Mudireddy M, Finke CM, Hanson CA, Ketterling RP, Szuber N, et al. Targeted next-generation sequencing in blast phase myeloproliferative neoplasms. Blood Adv. 2018;2:370–80.

37. Patel KP, Newberry KJ, Luthra R, Jabbour E, Pierce S, Cortes J, et al. Correlation of mutation profile and response in patients with myelofibrosis treated with ruxolitinib. Blood. 2015;126:790–7.

38. Rampal R, Ahn J, Abdel-Wahab O, Nahas M, Wang K, Lipson D, et al. Genomic and functional analysis of leukemic transformation of myeloproliferative neoplasms. Proc Natl Acad Sci U S A. 2014;111:E5401–10.

39. Bunda S, Burrell K, Heir P, Zeng L, Alamsahebpour A, Kano Y, et al. Inhibition of SHP2- mediated dephosphorylation of Ras suppresses oncogenesis. Nat Commun. 2015;6:8859.

40. Dance M, Montagner A, Salles JP, Yart A, Raynal P. The molecular functions of Shp2 in the Ras/Mitogen-activated protein kinase (ERK1/2) pathway. Cell Signal. 2008;20:453– 9.

41. Tiganis T, Bennett AM. Protein tyrosine phosphatase function: the substrate perspective. Biochem J. 2007;402:1–15.

42. Pandey G, Kuykendall AT, Reuther GW. JAK2 inhibitor persistence in MPN: uncovering a central role of ERK activation. Blood Cancer J. 2022;12:13.

43. Stivala S, Codilupi T, Brkic S, Baerenwaldt A, Ghosh N, Hao-Shen H, et al. Targeting compensatory MEK/ERK activation increases JAK inhibitor efficacy in myeloproliferative neoplasms. J Clin Invest. 2019;129:1596–611.

44. Winter PS, Sarosiek KA, Lin KH, Meggendorfer M, Schnittger S, Letai A, et al. RAS signaling promotes resistance to JAK inhibitors by suppressing BAD-mediated apoptosis. Sci Signal. 2014;7:ra122.

45. Brkic S, Stivala S, Santopolo A, Szybinski J, Jungius S, Passweg JR, et al. Dual targeting of JAK2 and ERK interferes with the myeloproliferative neoplasm clone and enhances therapeutic efficacy. Leukemia. 2021;35:2875–84.

46. Jayavelu AK, Schnoder TM, Perner F, Herzog C, Meiler A, Krishnamoorthy G, et al. Splicing factor YBX1 mediates persistence of JAK2-mutated neoplasms. Nature. 2020;588:157–63.

47. Nichols RJ, Haderk F, Stahlhut C, Schulze CJ, Hemmati G, Wildes D, et al. RAS nucleotide cycling underlies the SHP2 phosphatase dependence of mutant BRAF-, NF1- and RAS-driven cancers. Nat Cell Biol. 2018;20:1064–73.

48. LaRochelle JR, Fodor M, Vemulapalli V, Mohseni M, Wang P, Stams T, et al. Structural reorganization of SHP2 by oncogenic mutations and implications for oncoprotein resistance to allosteric inhibition. Nat Commun. 2018;9:4508.

49. Ember SW, Lambert QT, Berndt N, Gunawan S, Ayaz M, Tauro M, et al. Potent Dual BET Bromodomain-Kinase Inhibitors as Value-Added Multitargeted Chemical Probes and Cancer Therapeutics. Mol Cancer Ther. 2017;16:1054–67.

50. Lu X, Levine R, Tong W, Wernig G, Pikman Y, Zarnegar S, et al. Expression of a homodimeric type I cytokine receptor is required for JAK2V617F-mediated transformation. Proc Natl Acad Sci U S A. 2005;102:18962–7.

51. Brkic S, Meyer SC. Challenges and Perspectives for Therapeutic Targeting of Myeloproliferative Neoplasms. Hemasphere. 2021;5:e516.

52. Koppikar P, Bhagwat N, Kilpivaara O, Manshouri T, Adli M, Hricik T, et al. Heterodimeric JAK-STAT activation as a mechanism of persistence to JAK2 inhibitor therapy. Nature. 2012;489:155–9.

53. Kong T, Laranjeira ABA, Collins TB, De Togni ES, Wong AJ, Fulbright MC, et al. Pevonedistat targets malignant cells in myeloproliferative neoplasms in vitro and in vivo via NFkappaB pathway inhibition. Blood Adv. 2022;6:611–23.

54. Coltro G, Rotunno G, Mannelli L, Mannarelli C, Fiaccabrino S, Romagnoli S, et al. RAS/CBL mutations predict resistance to JAK inhibitors in myelofibrosis and are associated with poor prognostic features. Blood Adv. 2020;4:3677–87.

55. Kandarpa M, Robinson D, Wu YM, Qin T, Pettit K, Li Q, et al. Broad Next-Generation Integrated Sequencing of Myelofibrosis Identifies Disease-Specific and Age-Related Genomic Alterations. Clin Cancer Res. 2024;30:1972–83.

56. Maslah N, Kaci N, Roux B, Alexe G, Marie R, Pasquer H, et al. JAK2 inhibition mediates clonal selection of RAS pathway mutations in myeloproliferative neoplasms. Nat Commun. 2025;16:6270.

57. Mylonas E, Yoshida K, Frick M, Hoyer K, Christen F, Kaeda J, et al. Single-cell analysis based dissection of clonality in myelofibrosis. Nat Commun. 2020;11:73.

58. O’Sullivan JM, Taylor J, Gerds A, Buckley S, Harrison CN, Oh S, et al. RAS-pathway mutations are common in patients with ruxolitinib refractory/intolerant myelofibrosis: molecular analysis of the PAC203 cohort. Leukemia. 2023;37:2497–501.

59. Reynolds SB, Pettit K, Kandarpa M, Talpaz M, Li Q. Exploring the Molecular Landscape of Myelofibrosis, with a Focus on Ras and Mitogen-Activated Protein (MAP) Kinase Signaling. Cancers (Basel). 2023;15.

60. Santos FPS, Getta B, Masarova L, Famulare C, Schulman J, Datoguia TS, et al. Prognostic impact of RAS-pathway mutations in patients with myelofibrosis. Leukemia. 2020;34:799–810.

61. Fobare S, Kohlschmidt J, Ozer HG, Mrozek K, Nicolet D, Mims AS, et al. Molecular, clinical, and prognostic implications of PTPN11 mutations in acute myeloid leukemia. Blood Adv. 2022;6:1371–80.

62. Tartaglia M, Niemeyer CM, Fragale A, Song X, Buechner J, Jung A, et al. Somatic mutations in PTPN11 in juvenile myelomonocytic leukemia, myelodysplastic syndromes and acute myeloid leukemia. Nat Genet. 2003;34:148–50.

63. Baffert F, Regnier CH, De Pover A, Pissot-Soldermann C, Tavares GA, Blasco F, et al. Potent and selective inhibition of polycythemia by the quinoxaline JAK2 inhibitor NVP- BSK805. Mol Cancer Ther. 2010;9:1945–55.

64. Mazzacurati L, Collins RJ, Pandey G, Lambert-Showers QT, Amin NE, Zhang L, et al. The pan-PIM inhibitor INCB053914 displays potent synergy in combination with ruxolitinib in models of MPN. Blood Adv. 2019;3:3503–14.

65. Demaison C, Parsley K, Brouns G, Scherr M, Battmer K, Kinnon C, et al. High-level transduction and gene expression in hematopoietic repopulating cells using a human immunodeficiency [correction of imunodeficiency] virus type 1-based lentiviral vector containing an internal spleen focus forming virus promoter. Hum Gene Ther. 2002;13:803–13.

